# Feasibility of axon density metrics for brain asymmetry evaluation in the UK Biobank subsample

**DOI:** 10.1101/2020.02.25.965293

**Authors:** Ivan I. Maximov, Lars T. Westlye

## Abstract

Standard diffusion MRI model with intra- and extra-axonal water pools offers a set of microstructural parameters describing brain white matter architecture. However, a non-linearity of the general model and diffusion data contamination by noise and imaging artefacts make estimation of diffusion metrics challenging. In order to develop reproducible and reliable diffusion approaches and to avoid computational model degeneracy, one needs to devise additional theoretical assumptions allowing a stable numerical implementation. As a result, it is possible to estimate intra-axonal water fraction (AWF) representing one of the important structural parameters. AWF can be treated as an indirect measure of axon density and has a strong potential as useful clinical biomarker. A few diffusion approaches such as white matter tract integrity, neurite orientation dispersion and density imaging, and spherical mean technique, allow one to evaluate AWF in the frame of their theoretical assumptions. In the present study, we considered the compatibility of axon density metrics obtained from different diffusion models and the influence of the diffusion metric on a brain asymmetry estimation in UK Biobank sample consisting of 182 subjects. We found AWF derived from a spherical mean technique is the most statistically representative measure. As a result, we revealed that brain asymmetry indecies derived from intra-axonal water fraction weakly decrease along the lifespan, reducing the left-right hemisphere difference within increased age.

## Introduction

Diffusion MRI is a powerful non-invasive imaging technique allowing one to visualise and probe a living tissue at the micrometer scale. Random motions of water molecules affected by intra- and extra-cellular environments in white (WM) and grey matter can be caught and quantified by different diffusion encoding schemes. As a result, one can observe signal decay due to a spin phase decoherence originating from Brownian motion of water molecules (Jones, 2010). An important question arising in the interpretation of diffusion experimental data is how to connect the diffusion signal decay with underlying intra-voxel geometry and the organisation of complex living tissue (Novikov et al., 2018)? To answer to this question, many different diffusion models have been developed and applied to artificial systems (Fieremans and Lee, 2018), (Komlosh et al., 2017), (Vellmer et al., 2017a), animal models (Jespersen et al., 2019), (Ianuş et al., 2018), and in the human brain *in vivo* (Johansen-Berg and Behrens, 2014), (Jones, 2010).

An easy to use representation of diffusion imaging is a diffusion tensor imaging (DTI) (Basser et al., 1994), allowing one to introduce both a set of scalar metrics describing tissue integrity, such as fraction anisotropy (FA) or mean diffusivity, and WM connectivity using diffusion tensor eigenstates. Many attempts to fit the diffusion signal decay by different empirical functions have been made (Novikov et al., 2018) to suggest an accurate explanation of the brain microstructure and associated changes, for instance, axon losses or (de)myelination mechanisms (Johansen-Berg and Behrens, 2014). However, approaches based on signal fitting cannot explain an underlying tissue architecture and ongoing physiological processes (Novikov et al., 2018), (Novikov et al., 2019). In parallel, researchers have tried to model white and, partially, grey matter architecture by proposing a simplified representation of typical tissue compounds such as infinite cylinders, sticks and balls, impermeable spheres etc. For example, we can mention composite hindered and restricted model of diffusion (CHARMED) (Assaf and Basser, 2005), extended CHARMED model with introduced diameter distribution of restricted cylindrical axons (AxCalibre) (Assaf et al., 2008), neurite orientation distribution and density imaging (NODDI) (Zhang et al., 2012), white matter tract integrity (WMTI) (Fieremans et al., 2011), spherical mean techniques (SMT) (Kaden et al., 2016b), (Kaden et al., 2016a), restriction spectrum imaging (White et al., 2013), and more (Novikov et al., 2018). In turn, developed models have been validated using either artificial diffusion phantoms (Fieremans and Lee, 2018), (Komlosh et al., 2017), (Vellmer et al., 2017a) or ex vivo measurements (Ianuş et al., 2018), (Jespersen et al., 2019) including validation by a comparison with electron microscopy data (Lee et al., 2019). This has led to the formulation of a standard diffusion model based on decomposition of diffusion pools into intra- and extra-axonal water compartments (Novikov et al., 2018) (Novikov et al., 2019).

The standard diffusion model takes a non-linear approach and is associated with difficulties due to a flat solution landscape (Novikov et al., 2018), (Jespersen et al., 2019) and, as a result, a solution degeneracy. The use of the conventional Stejskal-Tanner pulse sequence with clinically feasible diffusion weightings (also known as *b*-values) does not allow one to resolve degeneration in diffusion imaging (Jelescu et al., 2015), (Jelescu et al., 2016). To avoid this problem, many biophysical models exploit additional assumptions simplifying the standard diffusion model and make it more practical. For example, equation linearisation (White et al., 2013), parameter fixation (Zhang et al., 2012), a priori known orientation distribution function (Tariq et al., 2016), have been used. Therefore, a clinical application of advanced diffusion models can be confined, in particular, in non-healthy tissue, due to the unknown distribution of pathological tissue parameters.

The healthy human brain possesses a strong quantitative microstructure variability in diffusion metrics, for instance, across age maturation or sex dimorphism (Smith et al., 2019), (Maximov et al., 2019), (Westlye et al., 2010). Large imaging databases, such as the UK Biobank (UKB), allow scholars to discover general brain patterns, exploiting imaging phenotypes that are accompanied by physiological, genetic and demographic data (Smith et al., 2019). UKB presents a great opportunity for brain research, in particular, using diffusion data (Elliott et al., 2018), (Smith et al., 2019). The chosen UKB diffusion protocol (Alfaro-Almagro et al., 2018) allows one to apply a set of biophysical models mentioned above. As a result, one can consider UKB as an excellent source for a statistical validation of diffusion models, in particular, in cases with unknown ground truth.

There is still a lack of direct diffusion metric comparisons with the same biophysical meaning (Jelescu et al., 2015), (Jelescu et al., 2016), (Coelho et al., 2019) using large subject cohorts. The intra-axonal water fraction which can be treated as an indirect axon density, has a unique representation of the WM organisation and, consequently, might play an important role as a powerful biomarker for further brain studies based on UKB: brain age gap evaluation (Smith et al., 2019), (Kaufmann et al., 2019), genome-wide association studies (Elliott et al., 2018), understanding of mental health disorders (Neilson et al., 2019). Thus, determining the accuracy and reproducibility of the diffusion-derived phenotypes is a crucial for further data analysis.

In the present work we aim to investigate whether axon density metrics derived from three popular diffusion models (WMTI, NODDI, and SMT) are correlated in healthy individuals. Due to the time consuming computations of NODDI parameters and impression of diffusion parameters fixation, it is important to understand whether NODDI metrics derived from different pipelines (Alfaro-Almagro et al., 2018), (Maximov et al., 2019) and algorithms (Zhang et al., 2012), (Daducci et al., 2015) are self-consistent and mutually correlated. To evaluate the efficiency of the axon density parameter as a sensitive biomarker, one can use brain structural asymmetry (Takao et al., 2011), (Takao et al., 2013) as a comparison criterion. Our results will yield a statistically representative and computationally easy-to-obtain biophysical model for UKB diffusion protocol which can be recruited in the future big-data analyses.

## Materials and Methods

### Participants and MRI data

In the present study we used 182 participants (age: min = 40.24; max = 70.11; mean = 54.70; std = 9.35 years), (sex: male = 90; female = 92). An accurate overview of the UKB data acquisition, protocol parameters, and image validation can be found in (Alfaro-Almagro et al., 2018), (Miller et al., 2016). Briefly, a conventional Stejskal-Tanner monopolar spin-echo echo-planar imaging (EPI) sequence was used with multiband factor 3, diffusion weightings were 1 and 2 ms/μm^2^ and 50 non-coplanar diffusion directions per each diffusion shell. All selected subjects were scanned at a single 3T Siemens Skyra scanner with a standard Siemens 32-channel head coil, in Cheadle, Manchester, UK. The spatial resolution was 2 mm^3^ isotropic, and 5 AP vs 3 PA images with b = 0 ms/μm^2^ were acquired. All diffusion data were post-processed using optimised diffusion pipeline (Maximov et al., 2019) consisting of 7 steps: noise correction (Veraart et al., 2016), Gibbs-ringing correction (Kellner et al., 2016), estimation of echo-planar imaging distortions, motion, eddy-current and susceptibility distortion corrections including outlier detection and reestimation (Andersson and Sotiropoulos, 2016), (Andersson et al., 2016), field non-uniformity correction (Tustison et al., 2010), spatial smoothing using *fslmaths* from FSL package (Smith et al., 2004) with the Gaussian kernel 1mm^3^, and diffusion metrics estimation (see below). A data quality was estimated by temporal signal-to-noise ratio (Roalf et al., 2016) for each *b*-shell. Original UKB data were estimated using UKB pipeline (Alfaro-Almagro et al., 2018) including susceptibility, eddy-current, and head motion corrections accompanied with slice outlier detection and replacement (Andersson et al., 2016), (Andersson and Sotiropoulos, 2016). The UKB scalar NODDI metrics were computed using Python implementation of the Accelerated Microstructure Imaging via Convex Optimisation (AMICO) algorithm (Daducci et al., 2015) of NODDI model (Zhang et al., 2012).

### Diffusion models

In order to derive axon density metrics based on intra-axonal water fraction we chose three biophysical models often used in clinical and research studies: WMTI, NODDI, and SMT, respectively. Below we briefly describe each approach.

### WMTI

In terms of the standard diffusion model, WMTI represents an intra-axonal space as a bundle of cylinders with effective radius equals to zero (Fieremans et al., 2011). The cylinders are impermeable, i.e., there are no water exchange between intra- and extra-axonal spaces. The extra-axonal space is described by anisotropic but still Gaussian diffusion. In order to keep the model simple, a few more assumptions have been made: that the intra-axonal space consists of mostly myelinated axons without any contribution from myelin due to fast relaxation rate across of typical diffusion times; at the same time in extra-axonal space the glial cells possess fast water exchange with extra-cellular matrix; both intra- and extra-axonal spaces are modelled by Gaussian diffusion tensors (Fieremans et al., 2011), (Jelescu et al., 2015). In order to avoid degeneration, it is assumed that diffusion inside of axons is slower than diffusion in extra-axonal matrix. Besides, WMTI parametrisation works in the case of a coherent axonal bundle with orientation dispersion below 30°. WMTI output consists of axonal water fraction (*AWF*), extra-axonal diffusivities: axial and radial components. The scalar metrics were estimated using original Matlab scripts (MathWorks, Natick, MA USA) from Veraart and colleagues (Veraart et al., 2013).

### NODDI

In comparison to WMTI, NODDI introduces three water compartments: intra- and extra-axonal spaces and isotropic water pool responsible for cerebrospinal fluid contamination (Zhang et al., 2012). The NODDI model assumes that the axonal bundle is coherent and axon orientation dispersion can be described by an axially symmetric function, such as Watson (Zhang et al., 2012) or Bingham (Tariq et al., 2016) functions. In turn, both intra- and extra-axonal diffusivities parallel to the bundle axis are fixed to plausible values (in the case of adults to 1.7 μm^2^/ms). The radial diffusivity in extra-axonal space is determined by the tortuosity model (Szafer et al., 1995): *D*^extra^ _┴_ = *D*^extra^ _||_ (1 – *f*_ic_), where *D*^extra^ are the extra-axonal radial (┴) and axial (||) diffusion coefficients, respectively; *f*_ic_ is the intra-axonal water fraction. Water diffusion in isotropic compartment is fixed to 3 μm^2^/ms. However, original NODDI approach computationally is time demanding. In order to accelerate an estimation of geometrical parameters, such as orientation dispersion and water fractions, Daducci and colleagues (Daducci et al., 2015) decomposed and linearised the problem using information about a bundle orientation from a DTI metric. This significantly reduced computation time per subject. The NODDI model output consists of intra-axonal water fraction (*icvf* for original NODDI metric, and *ICVF* for AMICO derived metric), isotropic water fraction, and orientation neurite dispersion. We estimated NODDI parameters using the Matlab scripts for original NODDI (Zhang et al., 2012) and for AMICO acceleration (Daducci et al., 2015).

### SMT

An estimation of orientation dispersion in the axon bundles is a complex theoretical problem (Novikov et al., 2019), in particular, using standard diffusion sequence protocols (Reisert et al., 2019). Recent achievements in isotropic diffusion weightings (Westin et al., 2016), (Jespersen et al., 2019), (Vellmer et al., 2017b) and double diffusion encoding (Henriques et al., 2019), (Shemesh et al., 2016) allowed one to avoid principle problems associated with standard model degeneration. In order to avoid a necessity to install a new pulse sequence on the clinical scanners one can recall a similar approach using a powder averaging technique (Kaden et al., 2016b) for both one and two compartment models (Kaden et al., 2016a). Nevertheless, an averaged signal still possesses a quite flat-fitting landscape, which might lead to degeneracy as in the case of NODDI (Jelescu et al., 2015), (Jelescu et al., 2016). As a result, Kaden et al. (Kaden et al., 2016a) increased the stability of the optimisation procedure by the following additional assumptions: diffusivity determines by the tortuosity model (Szafer et al., 1995), axial diffusivity in intra- and extra-axonal spaces are equal, and axons are presented as sticks, i.e. radial inta-axonal diffusion is equal to zero. In the present work we consider two compartment spherical mean technique (Kaden et al., 2016a) allowing one to extract an axon density metric. SMT output consists of intra-axonal water fraction (*intra*), intra-axonal diffusivity, and extra-axonal diffusivities: mean and radial components. We estimated SMT metrics using the original SMT code (https://github.com/ekaden/smt).

### Tract Based Spatial Statistics

In order to compare different diffusion metrics and approaches, we applied TBSS voxel-wise analysis (Smith et al., 2006). Initially, all volumes were aligned to the FMRI58_FA template, supplied by FSL (Smith et al., 2004), using a non-linear transformation implemented by FNIRT utility. Next, a mean FA images of all subjects was obtained and thinned in order to create mean FA skeleton. Afterwards, the subject’s FA values are projected onto the mean skeleton, by filling the skeleton with FA values from the nearest relevant tract centre. The skeleton-based analysis allows one to minimise confounding effects due to partial voluming and any residual misalignments originating from non-linear spatial transformations. Additionally, the TBSS derived skeleton is used for averaging of diffusion metrics over the skeleton.

We performed voxel-wise comparisons of NODDI scalar metrics obtained from two pipelines and different algorithm implementations using general linear model (GLM). For simplicity, we used individual level difference maps (*M*_A_−*M*_B_, where *M* is the scalar metric, and A/B are the algorithm or pipeline index) including age and sex as covariates. For all contrasts, statistical analysis was performed using permutation-based inference implemented in *randomise* with 5000 permutations. Threshold-free cluster enhancement (TFCE) was used (Smith and Nichols, 2009). Statistical *p*-value maps were thresholded at *p* < 0.05 corrected for multiple comparisons across space.

Brain asymmetry analysis was performed using the symmetrised TBSS skeleton produced by the FSL utility *tbss_sym*. The script generated the symmetric mean FA image and derived symmetric skeleton. Next, the difference maps between left-right hemispheres are voxel-wise evaluated for each diffusion metric using an appropriate design matrix and contrast files with age and sex as covariants by the *randomise* function with 5000 permutations. Statistical *p*-value maps were thresholded at *p* < 0.05 corrected for multiple comparisons as well.

### Linked independent component analysis

In order to model inter-subject variability across the diffusion metrics we performed data-driven decomposition based on linked independent component analysis (LICA) from the FSL package (Groves et al., 2011). The LICA approach is based on the conventional ICA technique, assuming that the signal presents a linear mixture of statistically independent spatial patterns. Along the optimisation, LICA iteratively searches maximally non-Gaussian patterns by subject weight updating. As a result, LICA components are characterised by spatial maps and subject individual weights. A model order was fixed by using cophenetic coefficient estimation (Ray et al., 2013).

The axonal water fraction metrics and FA maps were included in the LICA decomposition in order to evaluate the common and unique inter-subject variability across the six parameters taking into account differences in the pipeline and NODDI algorithm evaluations. Namely, FA from DKI, *AWF* from WMTI, *intra* from SMT, *icvf* from original NODDI and optimised pipeline, *ICVF* from AMICO NODDI and optimised pipeline, and *UICVF* from AMICO NODDI and UKB pipeline. Since FA values do not have direct meaning of axon density we repeated LICA analysis excluded FA maps as well. All LICA analyses were performed using TBSS skeletons with 1 mm^3^ resolution and 3000 iterations.

### Statistical analysis

Diffusion metrics and associated asymmetry indices (AI = [*M*_R_−*M*_L_]/[*M*_R_+*M*_L_], where R/L are the right or left values of metric *M*, respectively) were compared using general linear model (GLM) AI = *b*_0_ + *b*_1_ Age + *b*_2_ Sex. AI indices were averaged over the symmetrised TBSS skeleton after the estimation for the left and right hemispheres. The GLM fit was performed using Matlab function *lmfit*. The linear correlations between diffusion metrics and AI values were estimated using Matlab function *corr,* which produced a Pearson correlation coefficient.

## Results

Figure 1 shows the scatter plots of FA and estimated axonal water fractions from WMTI, SMT, and NODDI models. Diffusion metrics were averaged over the subject’s skeleton in accordance with the TBSS pipeline. The FA values demonstrated the lowest correlation coefficients among all diffusion metrics. The original NODDI metric (*icvf*) exhibits the same correlations (r = 0.97) with *ICVF* and *UICVF* values, in contrast to the lower correlation (r = 0.94) between the two NODDI AMICO metrics *ICVF* and *UICVF*. The *AWF* and *intra* (r = 0.97) metrics show a lower correlation (r = 0.92) with *ICVF* values. The histogram mode of axonal water fraction (0.39) for *AWF* values is lower than the modes (≈ 0.59) of all other metrics.

**Figure 1.**
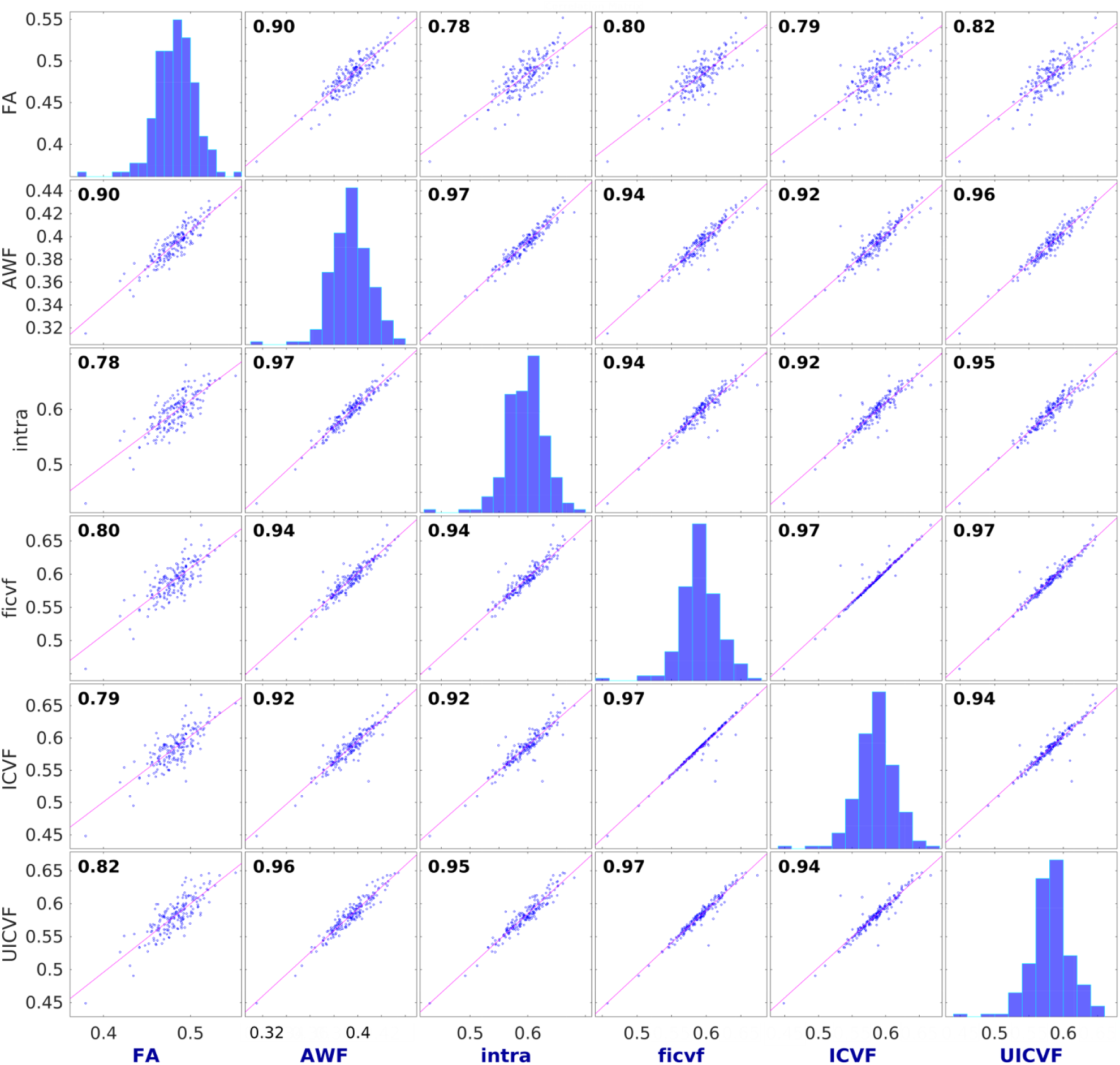
Correlation plots for FA and axonal water fractions obtained from WMTI (AWF), SMT (intra), and NODDI: original algorithm and optimised pipeline (ficvf); NODDI AMICO algorithm and optimised pipeline (ICVF); and NODDI AMICO algorithm and UKB pipeline (UICVF). All diffusion metrics were averaged over subject’s skeletons in accordance with TBSS pipeline.

In Figure 2, we present the results of LICA analysis using 20 independent components (IC) in two cases: with and without the FA metric. Fig. 2 shows the contribution of different diffusion metrics into each IC. The weights coefficient correlations of all subjects for 20 IC in both cases with and without FA metric are presented as well. As an example of common variation patterns, we present IC number 1 and 14. In the case of IC1 we find that all diffusion metrics play an important role in skeleton changes with the strongest contribution from *icfv* and *ICVF* metrics and lowest contribution from FA. In turn, in the case of IC14, all skeleton changes are determined by only *UICVF* contribution. As a result, we see that diffusion metrics *UICVF* and *ICVF* define ICs, where the diffusion contribution to IC is covered by almost one metric only.

**Figure 2.**
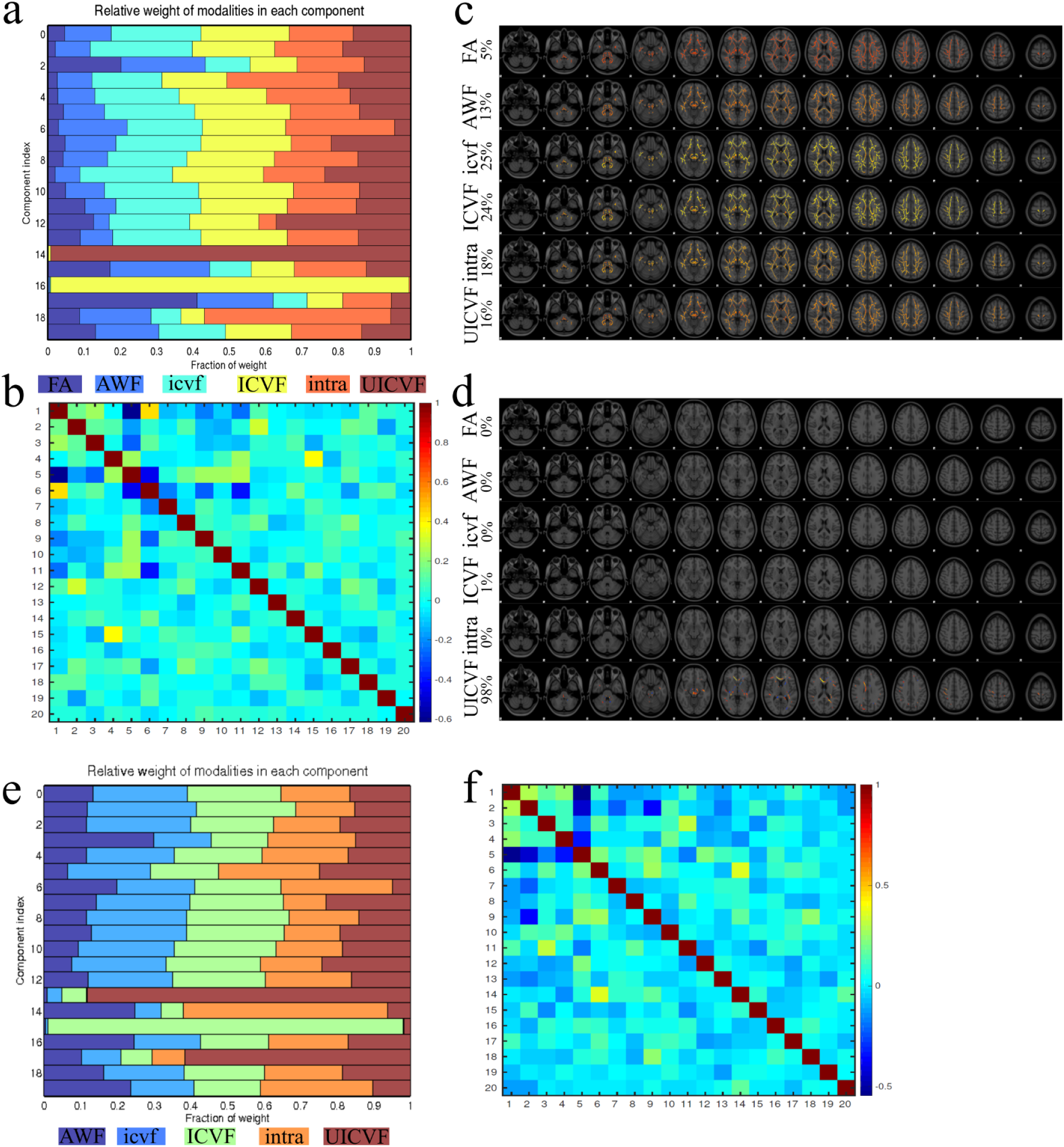
Results of LICA analysis based on diffusion metrics and TBSS skeleton with (a,c,d) and without (e,f) FA values. Number of independent components (IC) is equal to 20. a) contribution of diffusion metrics into IC; b) correlation map of weight coefficients for 20 IC; c) spatial patterns of common variance in the case of the first component; d) spatial patterns of common variance in the case of the 15^th^ component; e) contribution of diffusion metrics into IC; f) correlation map of weight coefficients for 20 IC.

In order to find spatial patterns with significant differences on the brain skeleton between the three NODDI approaches, we applied TBSS analysis for three NODDI-derived diffusion metrics (axon water fraction, isotropic water fraction, and bundle orientation dispersion). Figure 3 demonstrates the results of the TBSS analysis between two pairs of approaches: *a*) comparison between original NODDI and NODDI AMICO using the same optimised pipeline; *b*) comparison between optimised and UKB pipelines for NODDI AMICO algorithm implemented in Matlab (optimised pipeline) and in Python (UKB pipeline). Briefly, the difference between original NODDI and AMICO NODDI using the optimised pipeline covers a large part of the brain skeleton and presents a metric shift with underestimated values for AMICO NODDI. The metric shift appears in all NODDI-derived metrics. In turn, the difference for the same NODDI AMICO algorithm but with discrepancies in data preprocessing looks more dramatic and possesses complex skeleton patterns with under- and overestimated values. The differences are found in all NODDI metrics.

**Figure 3.**
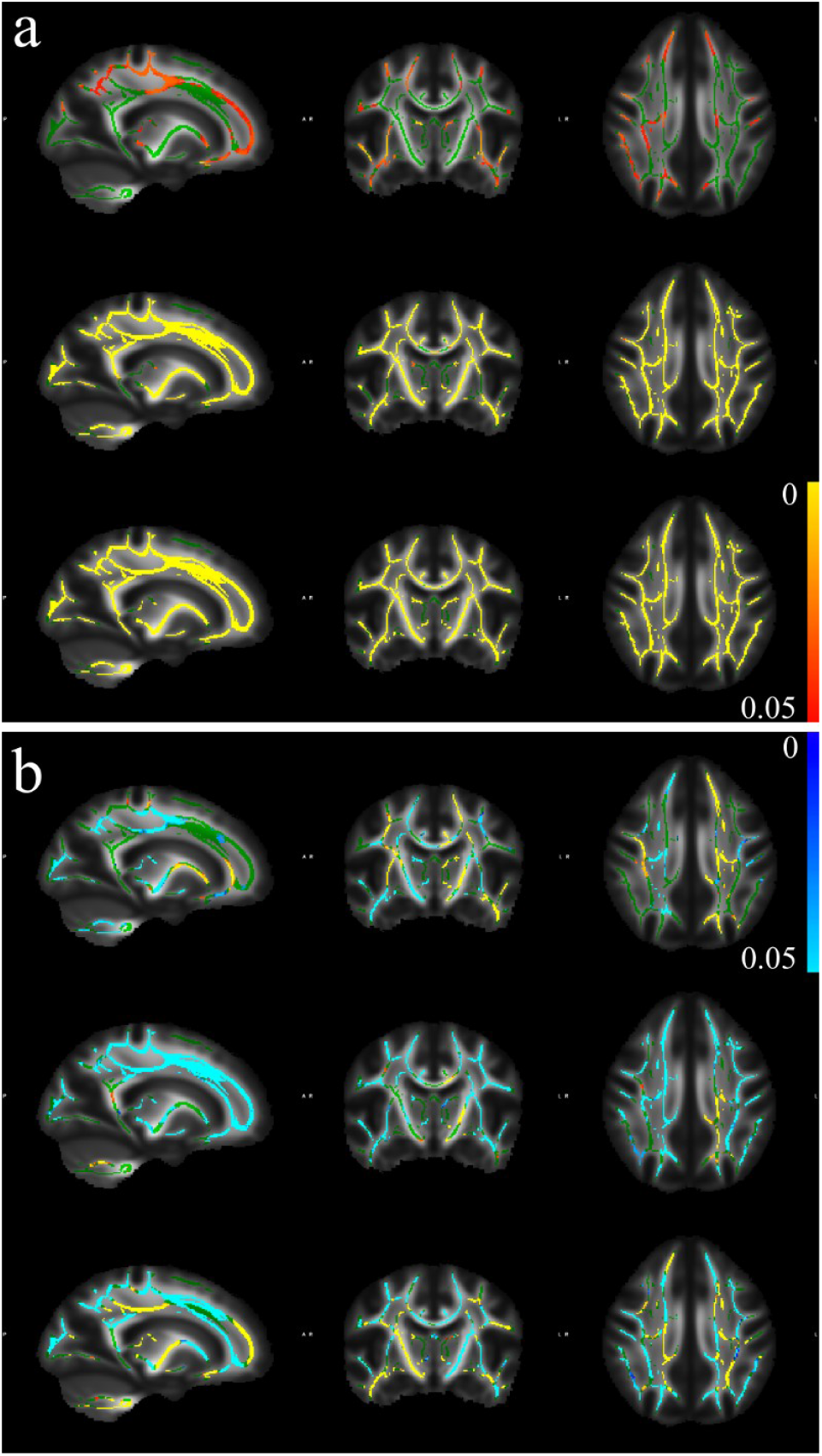
Voxelwise comparison of NODDI metrics using TBSS approach. **a**) comparison of original NODDI metrics vs NODDI AMICO estimations for axonal water fraction (top), isotropic water fraction (middle), and orientation distribution (bottom). The yellow-red colour marks the regions with values higher (p < 0.05) in original NODDI maps; **b**) comparison of NODDI AMICO metrics for optimised (Matlab scripts) and UKB (python scripts) pipelines. The top row images are cross-sections of axonal water fraction, the middle row is isotropic water fraction, and the bottom row is orientation distribution. The yellow-red colour marks the regions with values higher (p < 0.05) in optimised pipeline, the light blue-blue colour marks the regions with values lower (p < 0.05) in optimised pipeline. The skeleton is marked by the green colour.

Brain asymmetry was evaluated using the TBSS voxel-wise approach for symmetrised skeleton in accordance with the TBSS pipeline including age and sex as covariates. The results of the TBSS asymmetry analysis are presented in Figure 4. All diffusion metrics demonstrated regions with significant difference (p < 0.05), where the metrics are higher or lower in left hemisphere. In order to perform a pairwise comparison between statistically significant spatial patterns on the skeleton, we computed structural similarities between all image pairs using Matlab function *ssim* (Wang et al., 2004). The results are presented in Table 1. The structural similarities are estimated for two cases: the values from the left hemisphere are higher than in the right hemisphere, and in the opposite case, the values from the left hemisphere are lower than in the right hemisphere. These values are presented in the table cells over the main diagonal (the main diagonal marked by the red colour). The values below the main diagonal present a ratio between a number of common voxel and the total number of voxel with significant differences, i.e. R = A∩B/A, where A is the number of voxels in estimated skeleton region, and B is the number of voxels in skeleton region for a pair comparison. The comparison reveals that the spatial patterns with significant difference localised by *UICVF* metric demonstrate lower structural similarities among other diffusion metrics and a lower proportion of common skeleton voxels with significant difference. In turn, *icvf* and *ICVF* metrics demonstrate high level of proximity in both the structural similarities and number of the common voxels. Using data in Tab. 1 and the fact, that *ICVF* and *UICVF* metrics are quite unstable due to previous findings, we can evaluate a mean overlap between *AWF*, *intra* and *icvf* maps for higher and lower metrics in left hemisphere: *AWF* = 0.7913/0.5745; *intra* = 0.8254/0.6359; *icvf* = 0.8358/0.6098, respectively. These values allowed us to suggest the *intra* metric as having the highest overlapping rate among the diffusion metrics.

**Figure 4.**
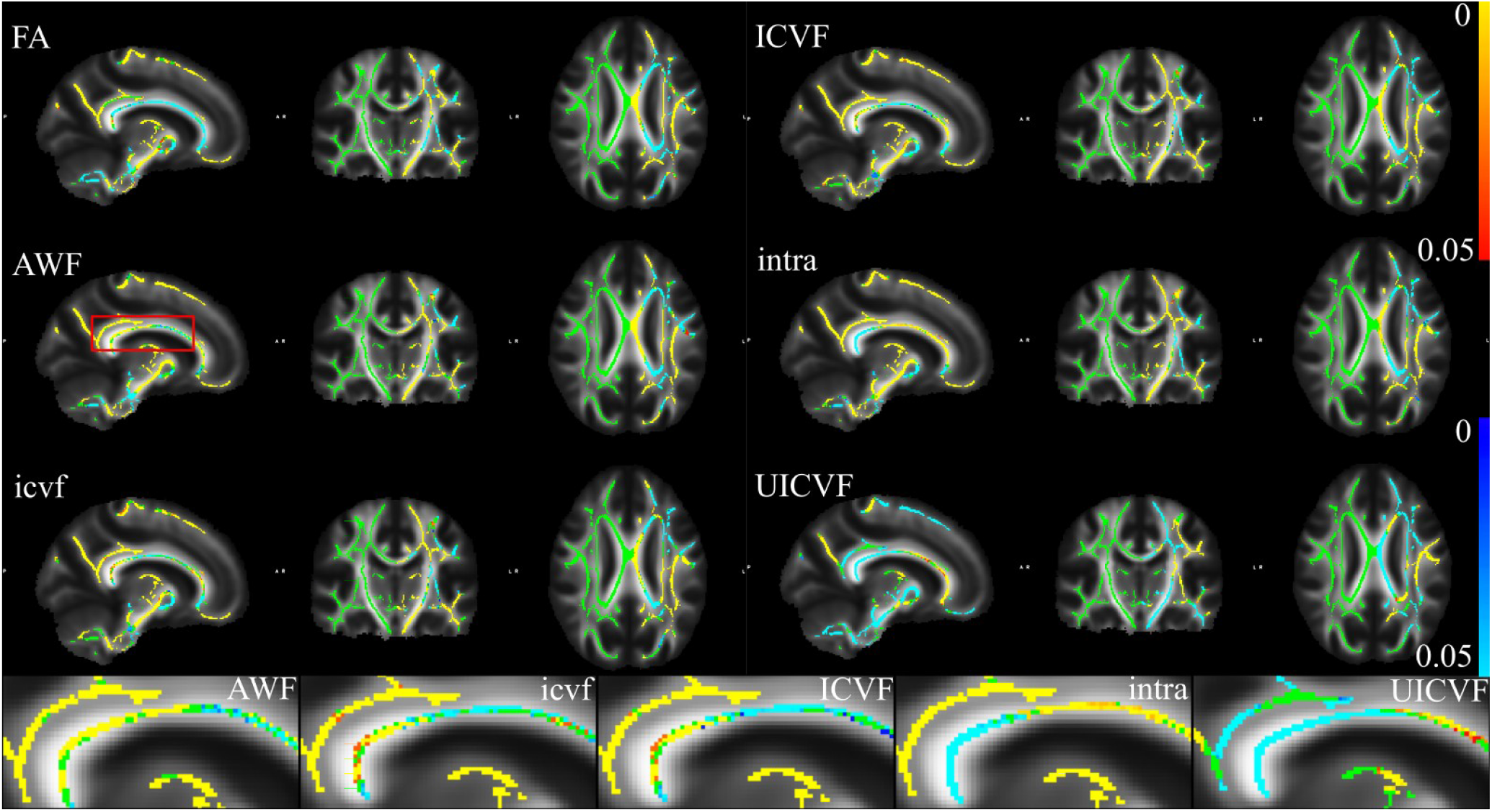
The result of TBSS analysis using the brain asymmetry feature. Diffusion metrics are represented by FA and axonal water fractions from different diffusion models. The yellow-red colour marks the regions with values higher (p < 0.05) in the left hemisphere, the light blue-blue colour marks the regions with values lower (p < 0.05) in the left hemisphere. The bottom row shows enlarged images localised by the red frame at AWF map. The symmetrised skeleton is marked by the green colour.

**Table 1.** A comparison of spatial patterns obtained by TBSS asymmetry analysis (p < 0.05). In upper diagonal cells we present pairwise SSIM estimations between the skeleton regions with significant differences. In bottom diagonal cells we present the pairwise ratio of voxel numbers N: (N_A_∩N_B_)/N_A_, where N_A,B_ is the number of voxel with significant difference from metrics A or B.

| **Values in the left hemisphere higher than in the right hemisphere** |  |  |  |  |  |
| --- | --- | --- | --- | --- | --- |
|  | **AWF** | **intra** | **icvf** | **ICVF** | **UICVF** |
| **AWF** | 1 | 0.9824 | 0.9819 | 0.9823 | 0.9608 |
| **intra** | 0.7809 | 1 | 0.9852 | 0.9860 | 0.9616 |
| **icvf** | 0.8017 | 0.8699 | 1 | 0.9972 | 0.9616 |
| **ICVF** | 0.8076 | 0.8797 | 0.9777 | 1 | 0.9614 |
| **UICVF** | 0.2068 | 0.1857 | 0.1850 | 0.1837 | 1 |
| **Values in the left hemisphere lower than in the right hemisphere** |  |  |  |  |  |
|  | **AWF** | **intra** | **icvf** | **ICVF** | **UICVF** |
| **AWF** | 1 | 0.9808 | 0.9804 | 0.9808 | 0.9605 |
| **intra** | 0.6006 | 1 | 0.9849 | 0.9857 | 0.9616 |
| **icvf** | 0.5483 | 0.6712 | 1 | 0.9968 | 0.9605 |
| **ICVF** | 0.5619 | 0.6956 | 0.9718 | 1 | 0.9604 |
| **UICVF** | 0.4151 | 0.4505 | 0.3530 | 0.3576 | 1 |

The scatterplots of diffusion metrics and derived AI values are presented in Figure 5. In order to evaluate a possible correlation between the diffusion metrics and their derived AI values we performed a linear regression with estimation of the Pearson correlation coefficients. The axon water fractions derived from the NODDI models demonstrated contradictory results with a positive correlation for AI vs *ICVF* values and negative correlations for AI vs (*icvf*, *UICVF*) values. Of note, in both cases the correlation coefficients were close to zero. In turn, the correlation between AI vs (FA, *AWF*, *intra*) metrics demonstrate a weak linear relationship. Interestingly, the AI vs FA scatter plot yields a simple rule for FA metrics and AI values: higher anisotropy tends to lower negative AI factor. The AI vs (*AWF*, *intra*) scatter plots show an opposite effect: the higher axon water fraction tends to higher positive AI values. The mode of AI histogram for *UICVF* locates in negative range (−0.01), when the modes for all other metrics are located in positive range. Thus, *UICVF* derived AI values demonstrate higher diffusion metric in the left hemisphere.

**Figure 5.**
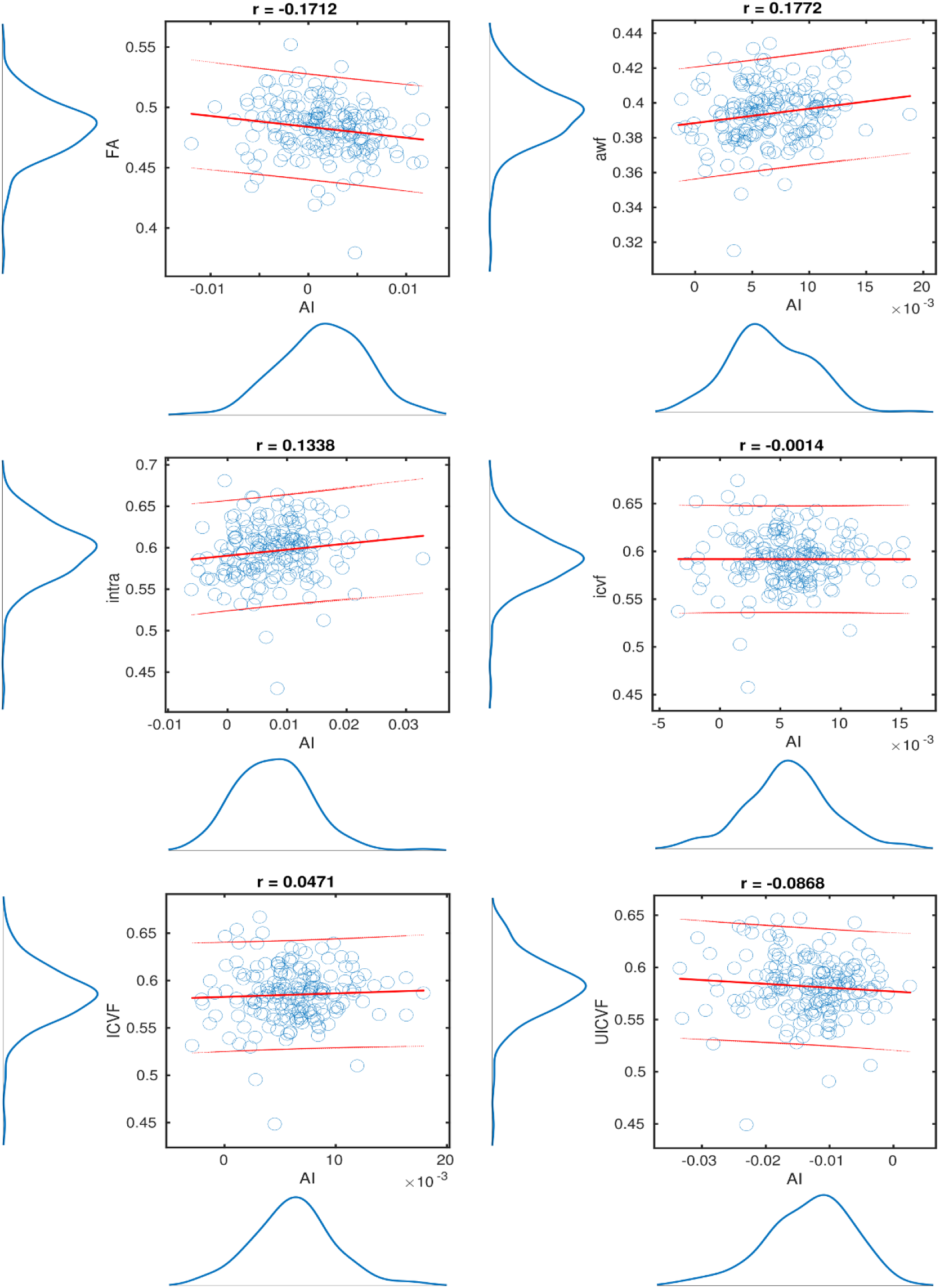
The result of linear regression of diffusion metrics and their derived AI. The red lines are linear regression fit and intervals of confidence (95%). The Pearson correlation coefficients are presented on the top of each correlation plot.

Finally, we estimated an influence of subject ages on the AI dependence using GLM. For this purpose, we fitted GLM with the sex as a covariance. The results are presented in Figure 6. In brief, the axon density metrics demonstrate a self-consistent behaviour by decreasing the AI along the subject years. In turn, the *b*_0_ intercept of the GLM demonstrated moderate variation among different diffusion models, in particular, for *UICVF*, which is negative. Interestingly, FA derived AI values weakly grow as age increases. The results of GLM fit such as intercept/slope and root mean squared error (RMSE) and R-squared values are summarised in Table 2. The highest R-squared and GLM slope values were found for *intra* and *UICVF* metrics.

**Figure 6.**
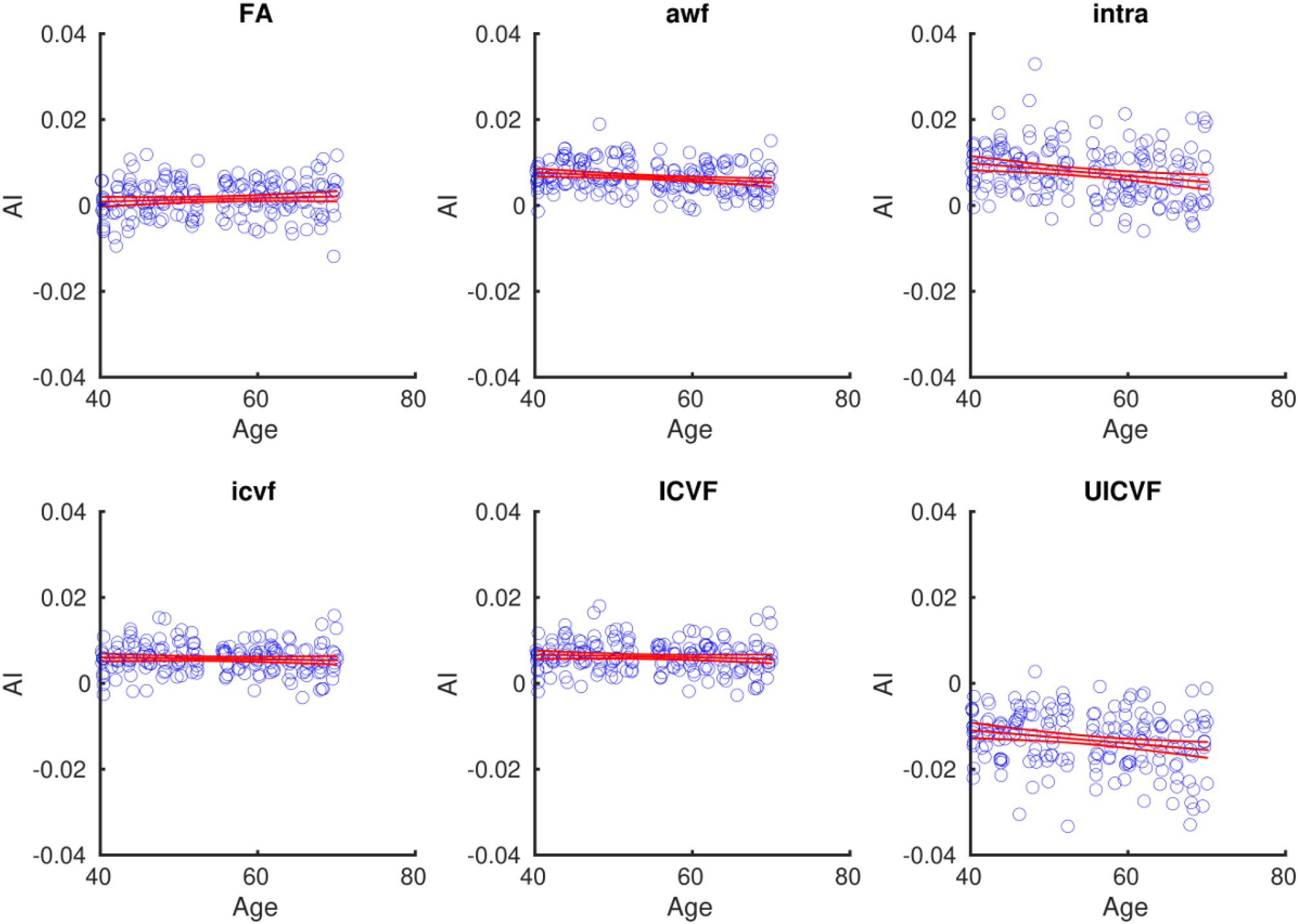
The results of GLM fit of age and AI dependences. The parameters of GLM fits are presented in Tab. 2. The red lines are linear regression fit and interval of confidence (95%).

**Table 2.** Parameters of regression fits for AI dependence on age using different diffusion metrics (see Fig. 6). The GLM is AI = *b*_0_ + *b*_1_ Age + *b*_2_ Sex. RMSE is the root mean squared error, R-squared is the coefficient of determination.

|  | **Intercept, $b_0$** | **Slope, $b_1$** | **RMSE** | **R-squared** |
| --- | --- | --- | --- | --- |
| **FA** | $-7.6880 \cdot 10^{-4}$ | $4.1383 \cdot 10^{-5}$ | 0.0042 | 0.0207 |
| **AWF** | 0.0108 | $-7.7402 \cdot 10^{-5}$ | 0.0035 | 0.0427 |
| **intra** | 0.0157 | $-1.4689 \cdot 10^{-4}$ | 0.0060 | 0.0512 |
| **icvf** | 0.0069 | $-2.2013 \cdot 10^{-5}$ | 0.0035 | 0.0036 |
| **ICVF** | 0.0080 | $-3.2509 \cdot 10^{-5}$ | 0.0036 | 0.0071 |
| **UICVF** | -0.0049 | $-1.5223 \cdot 10^{-4}$ | 0.0066 | 0.0504 |

## Discussion

Diffusion MRI is a sensitive tool for the estimation of the brain microstructure and its organisation, particularly in WM. The standard diffusion model (Novikov et al., 2019) is limited by non-linearity, numerical instability and being prone to image distortions. The proposed comparison of intra-axonal water fraction obtained from WMTI, NODDI, and SMT demonstrated that even very high correlations between these metrics does not guarantee a reproduction of the next statistical analysis. We found that NODDI metrics have significant voxel-wise differences depending on the estimation algorithm or preprocessing pipeline. In particular, the pipeline can significantly influence the analysis and leads to unreproducible results. LICA analysis demonstrated that the NODDI AMICO algorithm might seriously affect the diffusion metrics introducing algorithm specific variances and an estimation of ICs. The diffusion derived metrics such as asymmetry index, allowed us to suggest the SMT model as more statistically feasible using age-asymmetry association and diffusion-asymmetry distribution.

In the standard diffusion model (Novikov et al., 2018) the intra-axonal water fraction allows one to estimate indirect axon density and presents an important imaging phenotype. However, this parameter can be estimated using different theoretical assumptions in order to make the computations more reliable and stable in the frame of diffusion model. Thus, the same biophysical feature can be estimated from different diffusion models such as WMTI, NODDI, and SMT. In turn, even the same diffusion approach can have variability in numerical implementation such as original NODDI and NODDI AMICO. Does this approach variability produce self-consistent diffusion metrics? The skeleton averaged diffusion metrics demonstrated very high mutual correlations (see Fig. 1). However, even in this case, the correlation coefficients have variance depending on the diffusion model, for example, original NODDI metrics correlates with both NODDI AMICO metrics with r = 0.97. In turn, the same NODDI AMICO algorithm with differences in the pipeline exhibited a smaller correlation coefficient (r = 0.94). The axon density distribution also demonstrated the difference, since all model modes lie in close range of ~ 0.59, excepting WMTI with mode = 0.39.

LICA analysis allowed us to decompose a signal as a linear mixture of independent components. In the case of conventional analysis with multiple imaging modalities, it helps to specify spatial patterns with common variations and to estimate a contribution of each modality. Along decomposition of the same biophysical parameter, we expected to see a combination of all axon density metrics with a close percentage ratio. In the case of main independent components this hypothesis is realised (see Fig.2a,e). However, there are two diffusion models which suppressed all other contributions, namely, *ICVF* and *UICVF*, for higher order ICs. This kind of behaviour might be an indicator of unique features, not covered by other diffusion models. However, the original NODDI estimation did not recognise such a feature. It allows us to conclude that NODDI AMICO metrics demonstrated not specific behaviour of axonal water fraction. The spatial patterns defined by *UICVF* or *ICVF* are not as large as the results from main ICs (see Fig 2c,d), nevertheless, they might create spurious findings in the statistical analysis.

The LICA findings indicate that original NODDI and AMICO NODDI might have some serious differences based on numerical implementation of the same diffusion model. The voxel-wise TBSS analysis revealed that the significant differences are more dramatic, i.e. a use of the same post-processing pipeline created a value shift for NODDI AMICO metrics (see Fig. 3a). Notably, that for isotropic water faction and orientation dispersion metrics the value shift covered almost all skeleton regions. At the same time, the spatial patterns for intra-axonal water fraction have their own variability. Surprisingly, the optimised pipeline (Maximov et al., 2019) demonstrated a greater impact on the skeleton patterns with over- and under-estimated diffusion metrics (see Fig. 3b). It emphasises the need in accurately harmonised data before analysis, in particular, in the noise correction as an important step in post-processing pipeline and carefully to consider a numerical implementation of the used models (David et al., 2019), (Maximov et al., 2015).

Although the diffusion metrics reflect important microstructure brain features on their own, it can be used for a description of macroscale brain architecture such as brain asymmetry (Duboc et al., 2015). Brain asymmetry has potential as a useful biomarker (Zhong et al., 2016), in particular, in the case of mental disorders (Joo et al., 2018), (Wei et al., 2018), where the brain peculiarity is difficult to localise compared to the healthy brains. The estimated AI values based on different diffusion models and conventional FA using symmetrised TBSS skeleton revealed a moderate variability in the results, in particular, for *UICVF* data (see Fig. 4). Notably, that a coincidence of *icvf* and *ICVF* results is very high (over 97%). Nevertheless, the results are dependent on the diffusion metrics and a test design (higher or lower values in the left hemisphere).

Since AI values are derived from the symmetrised skeleton, it is quite interesting to verify their correlation with original diffusion metrics derived from the mean skeleton. The results of estimated correlations are presented in Fig. 5. We found that the correlation coefficients for *icvf*, *ICVF*, and *UICVF* scatter plots are negligibly low, in contrast to a weak relationship for FA, *AWF*, and *intra* metrics. Interestingly, that *icvf* and *UICVF* metrics exhibited the negative coefficients. The linear correlation between diffusion metric and AI values allowed us to assume that AI can be used as a complementary biomarker for age microstructure dependence.

The GLM fits of AI dependence on the age demonstrated self-consistent results for all axon density estimators. The regression parameters, summarised in Tab. 2, did not reveal significant differences between metrics. Nevertheless, the intercept of *UICVF*-age fit is negative in contrast to all other values. Surprisingly, *intra* values exhibited the higher RMSE and R-squared values in the regression, comparing to other modalities. The low regression slope for FA-age dependence partially reproduces the earlier published results (Takao et al., 2011). The AI behaviour demonstrated a presence of weak relationship between the brain asymmetry and subject’s age. In order to validate this finding and to increase the statistical power of this dependence, we plan to perform a more accurate analysis of the brain asymmetry using a whole UKB data with regional parcellation in the future (Maximov et al., 2020).

## Conclusion

In the present study, we considered the variability of one biophysical parameter of brain white matter derived from three diffusion approaches: WMTI, SMT, and NODDI. Our analysis suggests that axon density metrics based on NODDI approaches do not provide reliable quantities and depend on the numeric algorithm and post-processing pipeline. To estimate axonal water fraction, we recommend the use of the SMT model, which can be complementarily verified by *AWF* maps from WMTI. We found that the brain asymmetry measured by AI values derived from axon density is a useful and sensitive biomarker and can be used as an additional structural parameter in clinical and research studies. AI weakly changes along the lifespan and might be an additional covariant in the regression models.

## Acknowledgement

This work was funded by the Research Council of Norway (249795). This research has been conducted using the UK Biobank under Application 27412. Authors thank Dr. Daniel Quintana for proofread of the manuscript.

